# Enhancer-driven random gene overexpression (ERGO): a method to study gene function in Chlamydomonas

**DOI:** 10.1101/2025.09.05.674469

**Authors:** Yuliia Lihanova, Rory J. Craig, Mia Pribbernow, Reimund Goss, Severin Sasso

**Affiliations:** Institute of Biology, Department of Plant Physiology, Leipzig University, Leipzig, Germany; School of BioSciences, University of Melbourne, Parkville, VIC 3010, Australia

**Author notes:** Corresponding authors: Severin Sasso,; Yuliia Lihanova.

**Keywords:** *cis*-regulatory element/module (CRE/CRM), enhancer, activation tagging, gene overexpression/upregulation, gain-of-function, gene knockout/knockdown, loss-of-function, yellow-in-the-dark background, carotenoid metabolism, CRISPR/Cas9, leucine-rich repeat, F-box, forward genetics, green algae

## Abstract

Gene overexpression can be used to study gene function and is more suitable to characterize essential and redundant genes than gene knockout. A forward genetic approach based on random gene overexpression, also known as *activation tagging*, was previously used to study gene function in angiosperms. However, such an approach has never been applied to algae. Here, we present enhancer-driven random gene overexpression (ERGO), a forward genetic screen that we utilized to study genes involved in carotenoid metabolism in the green alga *Chlamydomonas reinhardtii*. We generated a library of over 33,000 insertional mutants in a yellow-in-the-dark background strain, which is incapable of producing chlorophyll in the dark. Each mutant contained a randomly inserted enhancer, *E*_*hist cons*_, capable of activating gene expression in the *C. reinhardtii* nuclear genome. After visually screening the mutant colonies for a color change from yellow to orange, we isolated a mutant with increased carotenoid content and remarkable resistance to high-light stress. RNA-seq data analysis revealed substantial upregulation of a gene, that we name *CMRP1*, encoding a putative F-box protein. CRISPR-mediated knockout of this gene resulted in decreased carotenoid concentrations, confirming that *CMRP1* is involved in the regulation of carotenoid metabolism. Our study shows that a gene overexpression screen can be successfully adapted to *C. reinhardtii* and potentially other plants and algae, thereby expanding the palette of genetic tools to study gene function.

## Introduction

Elucidation of gene function has been a major focus of biology for decades. Therefore, it is not surprising that a wide range of methods are available for studying gene function in different model organisms (Carpenter and Sabatini, 2004; Przybyla and Gilbert, 2022). These methods are based on two major gene modification techniques: loss of function and gain of function. Loss-of-function approaches, which completely prevent (knockout) or decrease (knockdown) gene expression, have been widely used to study gene function in eukaryotes in genome-wide functional screens (Przybyla and Gilbert, 2022). Over the years, extensive libraries of knockout mutants have been generated and many of them are commercially available (e.g. O’Malley and Ecker 2010; Lunardon et al., 2024). In contrast to loss of function, gain of function is based on the promoter- or enhancer-mediated increase in gene expression (Gou and Li, 2012). A main advantage of a gain-of-function over a loss-of-function approach is the possibility to study essential and redundant genes. Knocking out an essential gene will kill the organism, while a loss of function of only one of the redundant genes may not affect the phenotype due to the similar function and interchangeability of redundant genes (Nowak et al., 1997; Kuzmin et al., 2022). In such cases, gene overexpression represents a viable alternative to loss-of-function approaches.

Although gain of function has not been as widely used as loss of function, some studies in angiosperms successfully utilized an overexpression-based approach, called *activation tagging*. The concept of activation tagging was developed by Richard Walden’s group in the 1990s (Hayashi et al., 1992). In their pioneering approach, they designed a T-DNA-based transformation cassette with four enhancer copies from the cauliflower mosaic virus (CaMV) 35S gene and generated a library of insertional mutants with randomly overexpressed genes. The library was then screened to identify mutants with phenotypic changes. As a result, a gene involved in the auxin-independent growth of *Nicotiana tabacum* was identified (Hayashi et al., 1992). The activation tagging strategy developed in the Walden laboratory was later applied to other plant species, such as *Arabidopsis thaliana* (Robinson et al., 2009; Qin et al., 2022), *Oryza sativa* (Hsing et al., 2007; Gandikota et al., 2024), *Lotus japonicus* (Imaizumi et al., 2005), *Lycopersicon esculentum* (Mathews et al., 2003), *Populus tremula* (Busov et al., 2011), and *Petunia hybrida* (Zubko et al., 2002). Among the overexpressed genes, whose functions were characterized in detail, were genes involved in anthocyanin biosynthesis (Borevitz et al., 2000; Mathews et al., 2003), phytohormone metabolism (Zubko et al., 2002; Hsing et al., 2007), plant-pathogen interactions (Aboul-Soud et al., 2009), and the osmotic stress response (Koiwa et al., 2006).

While activation tagging has been successfully used in angiosperms, this method has not been applied to algae. In this study, we demonstrate how gene overexpression can be used in a forward genetic approach to elucidate gene function in the unicellular green alga *Chlamydomonas reinhardtii*. This model species has been extensively used to study different cellular processes, such as phototaxis, cell division, cell motility, photosynthesis, and stress responses, and is offering a wide range of genetic tools and extensive genomic resources (Sasso et al., 2018; Findinier and Grossman, 2023; Craig et al., 2023). Our strategy, called enhancer-driven random gene overexpression (ERGO), involved the generation and screening of a library of over 33,000 mutants. After phenotyping and RNA-seq data analysis of a promising mutant, we discovered a gene encoding a predicted F-box protein, whose overexpression caused an increase in carotenoid content. The high concentrations of carotenoids involved in protection against reactive oxygen species (ROS) led to a remarkable resistance of the mutant to high-light stress. Based on our first successful application of ERGO, we expect that this method will facilitate functional genomics research in green algae and beyond.

## Results

### A library of *C. reinhardtii* mutants with potentially overexpressed genes

To establish ERGO for the elucidation of gene function in *C. reinhardtii*, we used a forward genetic workflow (Fig. 1A). First, we generated a library of random insertional mutants by transformation with a cassette containing the enhancer *E*_*hist cons*_. This enhancer, which contains four copies of a highly conserved 8-bp motif and several other motifs located upstream of most H3/H4 histone genes, was shown to activate an endogenous promoter over a distance of at least 1.5 kb (Lihanova et al., 2024). To enable high-throughput screening of transformants for changes in carotenoid composition, we used the *C. reinhardtii* CC-3840 strain, known for its yellow-in-the-dark phenotype. This strain produces chlorophyll exclusively in the light using the light-dependent protochlorophyllide reductase (POR), whereas chlorophyll biosynthesis in the dark is prevented by a point mutation in the gene for the light-independent POR (Suzuki and Bauer 1992). This mutation results in a yellow color of the dark-grown cells due to the remaining carotenoids. The absence of the green chlorophylls *a* and *b*, which would have otherwise masked the yellow color of the carotenoids, allowed us to easily identify promising mutants by screening thousands of dark-grown mutant colonies for a color change from yellow to orange, brown, or red. One possible explanation for this color change was an increase in carotenoid content due to the *E*_*hist cons*_-induced gene overexpression. Thus, this screening strategy enables the discovery of genes involved in carotenoid metabolism and its regulation.

**Figure 1.**
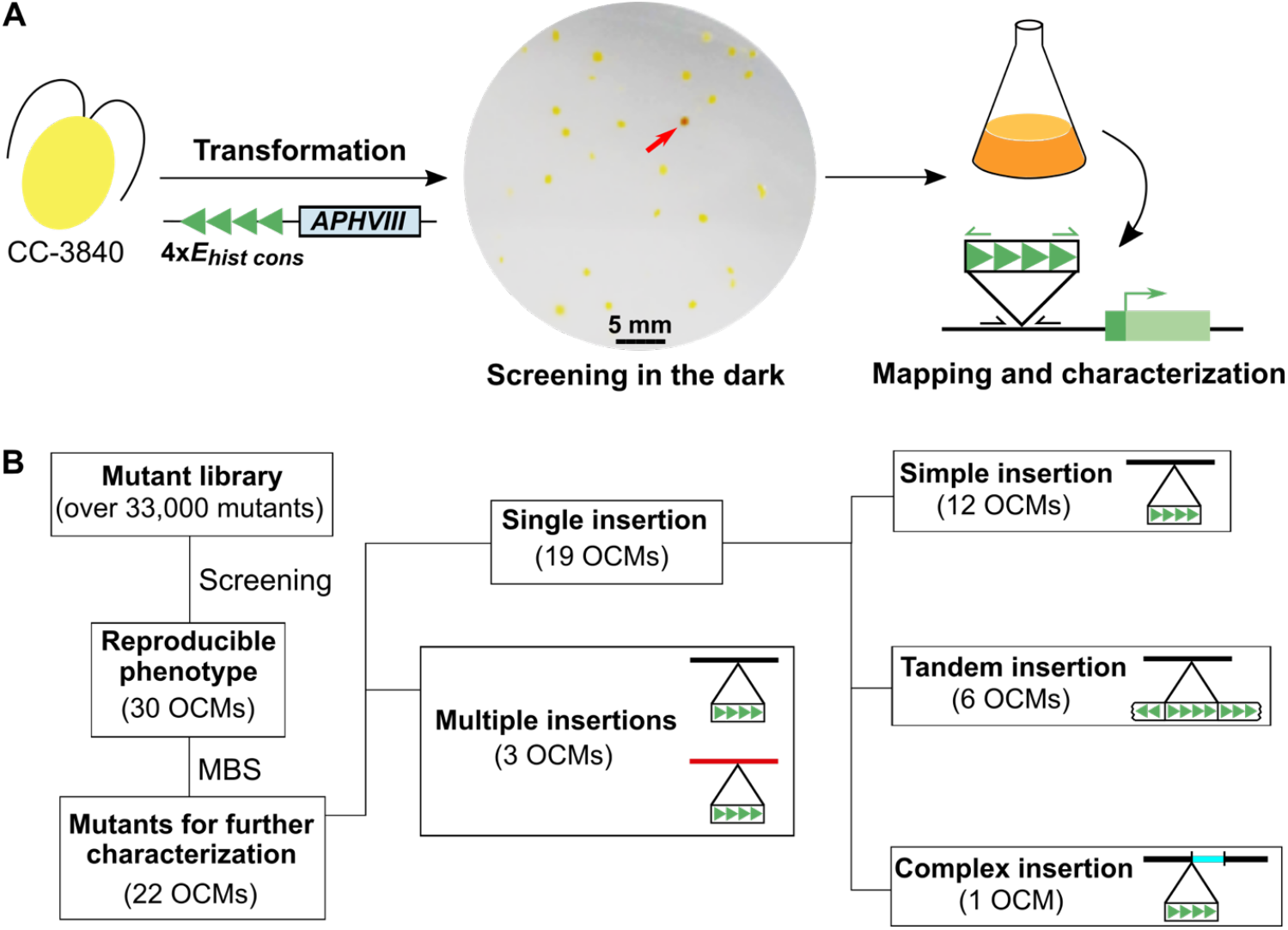
ERGO screen in *C. reinhardtii*. **(A)** ERGO workflow. *C. reinhardtii* strain CC-3840 with a yellow-in-the-dark phenotype was transformed by electroporation with a DNA cassette containing the enhancer *E*_*hist cons*_ with four copies of a conserved 8 bp-motif (four green triangles), and the *APHVIII* selection marker (Supplementary Fig. S1, S2). Transformants were screened for a visible color change on agar plates containing 15 µg ml^-1^ paromomycin in the dark. Mutants with a pronounced and reproducible phenotype were chosen for mapping of the *E*_*hist cons*_ insertion sites and further characterization. **(B)** Classification of orange-colored mutants (OCMs) based on the insertion type. The *E*_*hist cons*_ insertion sites were mapped in the nuclear genome of 30 OCMs by mapping-by-sequencing (MBS). In one OCM with a complex insertion, an exogenous 396 bp fragment of chloroplast DNA, co-inserted with the transformation cassette, is shown in blue. Enhancer insertions were sometimes accompanied by short indels or duplications of genomic DNA at the insertion site (Supplementary Data Set 1).

In total, we generated a library of over 33,000 mutants, each with a randomly integrated enhancer in its nuclear genome (Fig. 1B). Thirty mutants (0.1%) showed a reproducible color change from light yellow to various shades of orange in comparison with the background strain and were thus called orange-colored mutants (OCMs). These mutants were subjected to short-read Illumina or long-read PacBio whole-genome sequencing to map the *E*_*hist cons*_ insertion sites (Supplementary Data Set 1). As a result, we determined the *E*_*hist cons*_ insertion sites in 22 out of 30 OCMs. In eight OCMs, the insertion sites could not be mapped due to low coverage (4 OCMs), or the transformation cassette was truncated and did not contain *E*_*hist cons*_ (4 OCMs). Out of the remaining 22 OCMs, 19 OCMs had a single insertion at one locus, and three OCMs had multiple insertions at different loci. Mutants with single insertions were further classified depending on the complexity of the cassette insertion (Fig. 1B). We identified simple insertions of a complete transformation cassette, and tandem insertions that always consisted of truncated cassette fragments. One mutant had a complex insertion where the transformation cassette co-inserted with an exogenous fragment of chloroplast DNA, possibly originating from lysed cells present during electroporation (Zhang et al., 2014).

### The mutant OCM-43 can withstand high-light stress

In this study, we present a more detailed characterization of the mutant OCM-43. This mutant showed a reproducible color change to dark yellow compared to the light-yellow background strain CC-3840 (Fig. 2A). To explain this color difference, we quantified the concentrations of different carotenoids, including zeaxanthin, lutein, and β-carotene, which protect algal cells against the damaging effects of ROS (Wakao and Niyogi, 2021). Both in the dark and under continuous light of 50 µmol photons m^-2^ s^-1^, OCM-43 produced larger amounts of xanthophylls, such as violaxanthin, antheraxanthin, zeaxanthin, lutein, as well as β- and α-carotenes, when compared to CC-3840 (Fig. 2A). For most pigments, the absolute increases in concentrations between OCM-43 and CC-3840 were even more pronounced in the presence of light (Fig. 2A), when carotenoid biosynthesis is known to be upregulated (Bohne and Linden, 2002).

**Figure 2.**
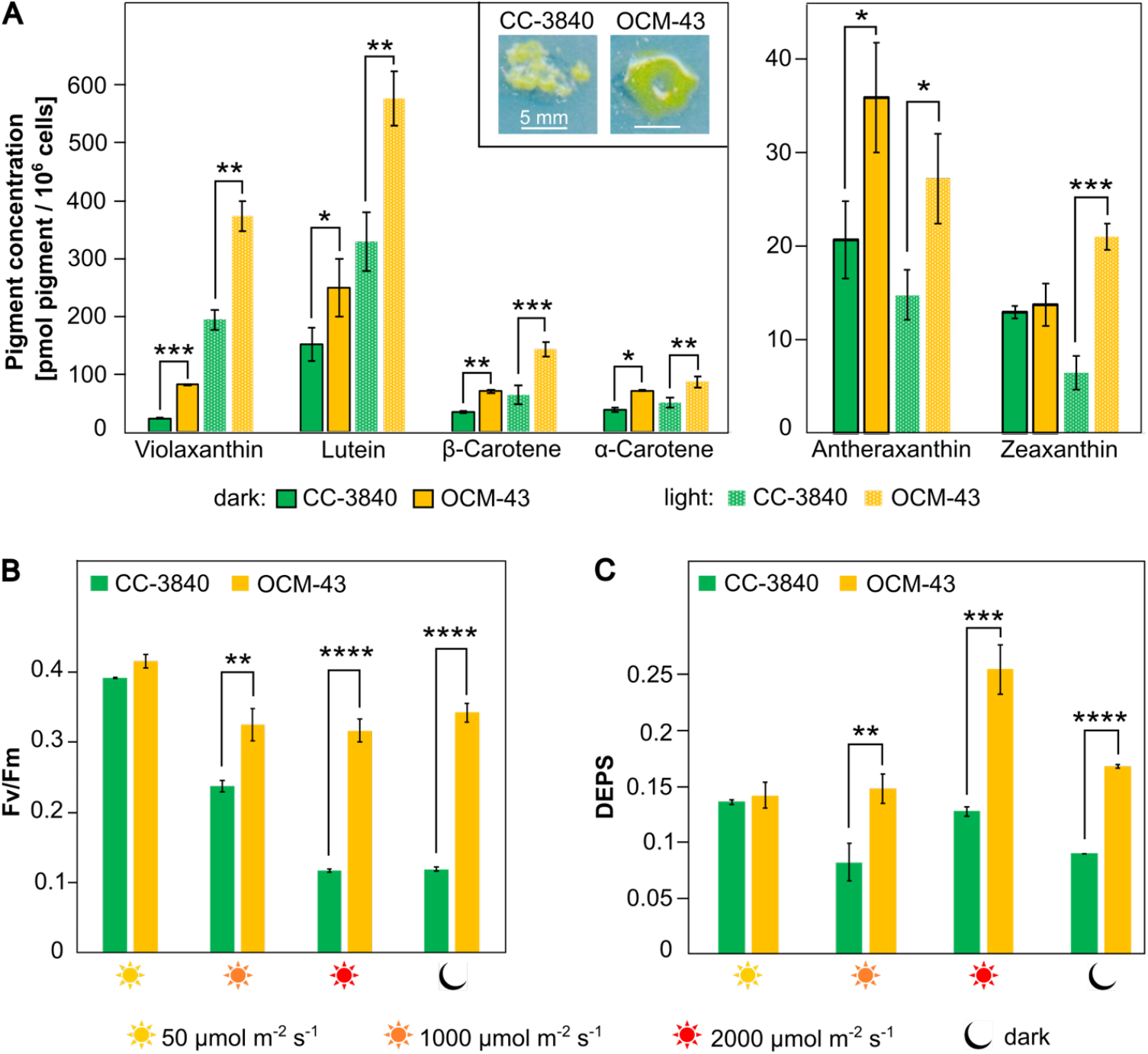
Phenotypic characterization of the mutant OCM-43. **(A)** Carotenoid concentrations in OCM-43 and CC-3840. Cultures were grown in TAP medium in the dark or under continuous illumination with a light intensity of 50 µmol photons m^-2^ s^-1^. Only carotenoids whose concentrations were significantly different between OCM-43 and CC-3840 are shown. The concentrations of the carotenoids neoxanthin and loroxanthin, and of chlorophylls *a* and *b* are shown in Supplementary Fig. S3. Inset: dark-yellow color of OCM-43 compared to the light-yellow background strain CC-3840 when grown on a TAP agar plate in the dark. **(B)** Quantum yield of PSII (Fv/Fm) in OCM-43 and CC-3840 under high-light stress. Cultures were grown in TP medium under a light intensity of 50 µmol photons m^-2^ s^-1^ and exposed to high light of 1000 µmol photons m^-2^ s^-1^ or 2000 µmol photons m^-2^ s^-1^ for 4 h. After 4 h at 2000 µmol photons m^-2^ s^-1^, a recovery phase of 1 h in the dark was included. **(C)** De-epoxidation state (DEPS) of the xanthophyll cycle pigments in OCM-43 and CC-3840. Cultivation conditions are described in (B). In (A), (B), and (C), mean values were compared with each other using an unpaired t-test. Statistically significant differences between the mean values are indicated as follows: * p ≤ 0.05; ** p ≤ 0.01; *** p ≤ 0.001; **** p ≤ 0.0001. Data analysis was based on three biological replicates with the values indicating the mean ± standard deviation.

To test whether the elevated carotenoid content facilitates the mutant’s survival under high-light conditions, we measured the quantum yield of photosystem II (PSII), Fv/Fm. This parameter has been widely used to characterize the photosynthetic activity of plants (Schreiber 2004). Cultures, grown under continuous light of 50 µmol photons m^-2^ s^-1^, were exposed to four hours of prolonged illumination with high light of 1000 µmol photons m^-2^ s^-1^ or 2000 µmol photons m^-2^ s^-1^. After exposure to 2000 µmol photons m^-2^ s^-1^, the cultures were incubated in the dark for one hour to test if Fv/Fm remained at low values, indicating sustained photoinhibition. After high-light illumination, CC-3840 showed low Fv/Fm values which did not recover during one hour of recovery phase in the dark (Fig. 2B). This result is in line with previous studies, where a severe inhibition of photosynthesis by high-light illumination, known as photoinhibition, has been observed in *C. reinhardtii* (Erickson et al., 2015). In contrast, OCM-43 showed higher Fv/Fm values after high-light exposure and a slight recovery of the PSII quantum yield during the dark period following illumination, arguing for a strongly reduced photoinhibition and thus an increased photoprotection of this mutant (Fig. 2B). In agreement with the enhanced high-light tolerance, OCM-43 showed a considerably higher de-epoxidation state of the xanthophyll cycle pigments (DEPS) after the illumination compared with the background strain (Fig. 2C). The DEPS describes to which extent violaxanthin has been de-epoxidized to the photoprotective pigments, antheraxanthin and zeaxanthin, during the operation of the xanthophyll cycle (Färber and Jahns, 1998). The increased concentrations of antheraxanthin and zeaxanthin, together with the increased amounts of the other photoprotective pigments, such as lutein and β-carotene, provide the basis for the increased photoprotection of OCM-43.

### Gene upregulation is a predominant response to the enhancer integration in OCM-43

Mapping-by-sequencing of OCM-43 revealed that two *E*_*hist cons*_ copies are inserted in a non-coding region on chromosome 17, which was additionally validated by Sanger sequencing (Supplementary Fig. S4). Therefore, *E*_*hist cons*_ may have activated proximal or distal genes on chromosome 17 (Fig. 3A). To identify differentially expressed genes, we analyzed the RNA-seq data of OCM-43 in comparison with CC-3840 (Supplementary Data Set 2). A principal component analysis demonstrated that the gene expression profile of the mutant was different from the profile of the background strain (Fig. 3B). Indeed, a total of 2855 genes out of 17,926 genes (15.9%) were differentially expressed in OCM-43 compared to CC-3840 (log_2_FC ≥ 1; *P*_*adj*_ ≤ 0.01), with the number of downregulated genes being slightly higher than that of the upregulated genes (Fig. 3C). However, when considering the genes with the most significant changes, most of them were upregulated with increases in transcript levels of up to 100-fold (Fig. 3D; Table 1). Seven of the top 20 upregulated genes belonged to the chloroplast genome and encoded subunits of RNA polymerase III, ribosomal proteins, and components of photosystems I and II (Fig. 3D; Table 1). Upon further inspection of the top 100 upregulated genes, 18 genes in total belonged to the chloroplast genome, and 25 genes from the nuclear genome encoded predicted plastid-targeted proteins (Supplementary Data Set 2). The higher concentrations of different chloroplast components, particularly PSI and PSII proteins, may provide the binding sites for the increased number of carotenoid molecules in OCM-43, thereby contributing to the enhanced photoprotection of the mutant.

**Table 1.** The top 20 differentially expressed genes in OCM-43 (sorted by *P*_*adj*_). Positive log_2_FC values indicate upregulation, while negative values indicate downregulation of gene expression in OCM-43 compared to CC-3840. Since enhancer *E*_*hist cons*_ was inserted on chromosome 17, the top 20 upregulated genes on this chromosome were analyzed in more detail. Functional annotations of the examined genes were retrieved from the plant genome database Phytozome (Goodstein et al., 2012). Identifiers of genes with an experimentally validated function are highlighted in bold. Analysis was based on the *C. reinhardtii* CC-4532 v6.1 genome (Craig et al., 2023). *P*_*adj*_, adjusted *P*-value; FC, fold change.

| Gene identifier | $P_{adj}$ | $\log_2FC$ | Functional annotation |
| --- | --- | --- | --- |
| **Differentially expressed genes, whole genome** |  |  |  |
| Cre17.g734805 | 2.32E-126 | 4.78 | Protein with a predicted leucine-rich repeat domain |
| **CreCp.g802335** | 1.32E-103 | 4.72 | RNA polymerase III, beta' chain |
| Cre12.g801441 | 2.27E-72 | 5.34 | - |
| Cre13.g602750 | 8.09E-63 | -3.72 | AP2-like ethylene-responsive transcription factor |
| Cre20.g751697 | 1.13E-60 | -5.49 | Sterol esterase / Triterpenol esterase |
| Cre07.g800869 | 5.69E-55 | 3.58 | ATP pyrophosphate-lyase |
| Cre12.g526240 | 1.41E-54 | 4.19 | Cytochrome c1 heme-lyase |
| Cre03.g143967 | 6.31E-54 | -3.83 | - |
| **CreCp.g802298** | 2.29E-52 | 3.36 | RNA polymerase III, alpha subunit |
| Cre01.g033500 | 1.58E-51 | 2.90 | Sphingomyelin phosphodiesterase |
| **Cre02.g146950** | 9.46E-51 | 2.73 | RNA lariat debranching enzyme |
| **CreCp.g802316** | 2.10E-50 | 3.26 | Heme-binding protein |
| Cre09.g400404 | 1.75E-49 | 5.98 | - |
| Cre12.g541400 | 1.73E-45 | -4.49 | Las17-binding protein actin regulator |
| **CreCp.g802306** | 9.56E-45 | 3.08 | RNA polymerase III, beta subunit I |
| Cre02.g095151 | 6.61E-44 | 6.66 | Sulfate-transporting ATPase |
| **Cre04.g222500** | 1.22E-42 | 2.46 | Bicaudal D (basal body proteome) |
| **CreCp.g802275** | 6.14E-42 | 3.04 | 30S ribosomal protein S19 |
| **CreCp.g802324** | 2.57E-40 | 2.97 | Mediator of the chloroplast protein import |
| **CreCp.g802265** | 7.20E-40 | 3.00 | Light-independent protochlorophyllide reductase, subunit B |
| **Upregulated genes, chromosome 17** |  |  |  |
| Cre17.g734805 | 2.32E-126 | 4.78 | Protein with a predicted leucine-rich repeat domain |
| Cre17.g745997 | 3.66E-26 | 2.80 | Threonine-specific protein kinase |
| Cre17.g729400 | 3.20E-16 | 3.68 | ATP pyrophosphate-lyase |
| Cre17.g747497 | 1.39E-14 | 1.94 | - |
| **Cre17.g721500** | 3.58E-13 | 1.29 | Granule-bound starch synthase IA |
| **Cre17.g740950** | 1.39E-12 | 2.90 | LHC-like protein |
| Cre17.g802142 | 1.14E-11 | 3.21 | Histone acetyltransferase |
| Cre17.g802076 | 1.73E-11 | 1.54 | Antifreeze protein, type I / Vacuolar protein sorting-associated protein 13 |
| Cre17.g719450 | 2.17E-11 | 1.34 | Threonine-specific protein kinase |
| Cre17.g802051 | 2.14E-10 | 1.89 | Transposase DNA-binding domain |
| **Cre17.g696500** | 1.13E-09 | 2.23 | Pherophorin-Chlamydomonas homolog 19 |
| Cre17.g712900 | 2.33E-09 | 1.57 | - |
| Cre17.g802114 | 1.28E-08 | 2.35 | Transposase DNA-binding domain |
| Cre17.g720850 | 2.38E-08 | 1.71 | - |
| Cre17.g704000 | 3.11E-08 | 2.58 | Polyvinyl-alcohol oxidase |
| Cre17.g729101 | 3.23E-08 | 4.09 | Pherophorin-Chlamydomonas homolog |
| Cre17.g716650 | 8.47E-08 | 1.18 | Ubiquitin-related domain |
| Cre17.g802063 | 9.53E-08 | 1.45 | - |
| Cre17.g802035 | 6.16E-07 | 1.68 | - |

**Figure 3.**
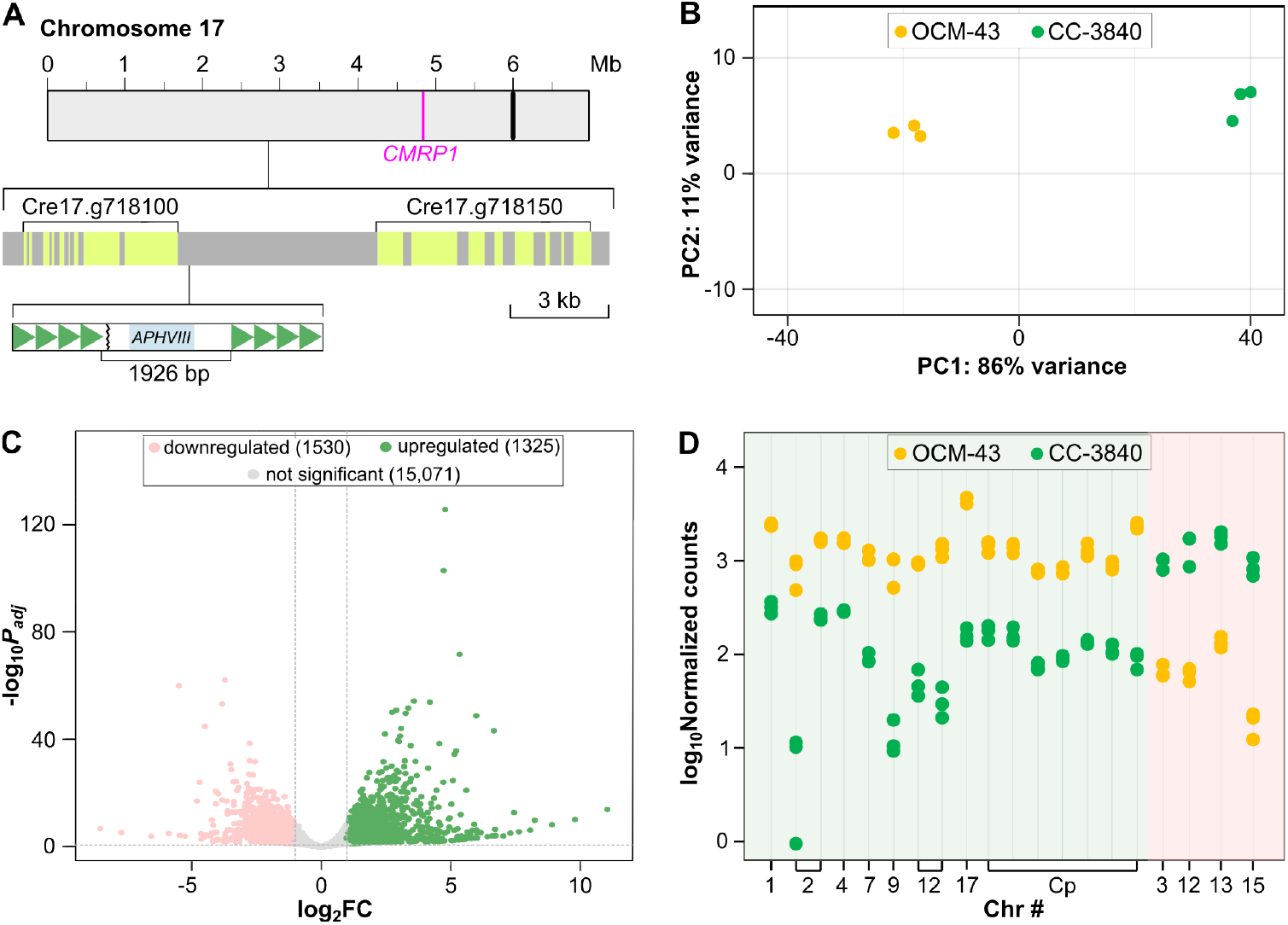
Insertion site and RNA-seq data analysis of mutant OCM-43. **(A)** Insertion site of *E*_*hist cons*_ in the OCM-43 nuclear genome. Two fragments of the transformation cassette are inserted in the non-coding region of chromosome 17, resulting in two copies of *E*_*hist cons*_ (dark-green triangles) separated by 1926 bp of the cassette sequence. Coding regions that correspond to the exons of nearby genes are shown in light green. The centromere of chromosome 17 is shown by a black vertical line. The magenta line indicates genomic location of the most significantly upregulated gene, *CMRP1* (Cre17.g734805). **(B)** Principal component (PC) analysis of the RNA-seq data of OCM-43 and the background strain CC-3840 grown under continuous low light of 15 µmol photons m^-2^ s^-1^. Dots represent three biological replicates. **(C)** Volcano plot of differentially expressed genes in OCM-43. The change in gene expression was considered statistically significant if |log_2_FC| was ≥ 1 and *P*_*adj*_ ≤ 0.01. FC, fold change; *P*_*adj*_, adjusted *P*-value. **(D)** Normalized count values for the top 20 differentially expressed genes in OCM-43. The top 20 genes were determined by sorting all genes from the lowest to the highest *P*_*adj*_ value. Three biological replicates are depicted for each gene of OCM-43 and CC-3840. Upregulated genes are shown on the pale green background, and downregulated genes are highlighted by pink background. Analyses were based on the *C. reinhardtii* CC-4532 v6.1 genome (Craig et al., 2023). Chr #, chromosome number; Cp, chloroplast genome.

Since *E*_*hist cons*_, which enhances gene expression in *cis* (Lihanova et al., 2024), was inserted into chromosome 17, we examined the upregulated genes on this chromosome in more detail. The genes flanking *E*_*hist cons*_ (Cre17.g718100, Cre17.g718150) were not differentially expressed in OCM-43 (Supplementary Data Set 2). The most significantly upregulated gene on chromosome 17, which we called *CMRP1* (for carotenoid metabolism-regulating protein; Cre17.g734805) showed the largest increase in expression of almost 30-fold and is located approximately 2 Mb away from the enhancer (Fig. 3A). This gene encodes a protein with a predicted leucine-rich repeat domain (Table 1). Notably, *CMRP1* was also the most significantly differentially expressed gene in the whole genome (Table 1), and we thus investigated more closely whether the upregulation of this gene is responsible for the carotenoid accumulation in OCM-43 (genotype-phenotype link).

### Discovery of a new gene involved in the regulation of carotenoid metabolism

The gene *CMRP1* encodes a protein of 2577 amino acids with two predicted leucine-rich repeat (LRR) domains (Fig. 4A). LRR domains have been implicated in protein-protein interactions and are often present in F-box proteins, key components of the ubiquitin-proteasome system (Zhang et al., 2018). Therefore, we searched for any additional domains in the CMRP1 amino acid sequence based on a multiple sequence alignment built from PSI-BLAST queries (Supplementary Data Set 3). As a result, we identified an F-box domain whose sequence is interrupted by an insert of 99 amino acids in CMRP1 (Fig. 4A), the presence of which complicated motif annotation. Based on the structural prediction, this insert folds into a loop, bringing the F-box helices in close proximity (Fig. 4A). Orthologs with an F-box domain, similarly interrupted by an insert, are also present in other volvocine algae (Fig. 4B; Supplementary Fig. S5). In addition, homology of the CMRP1 protein could be traced throughout green algae and to land plants, although the insert interrupting the F-box motif is absent from the respective sequences (Supplementary Fig. S5; Supplementary Data Sets 3, 4).

**Figure 4.**
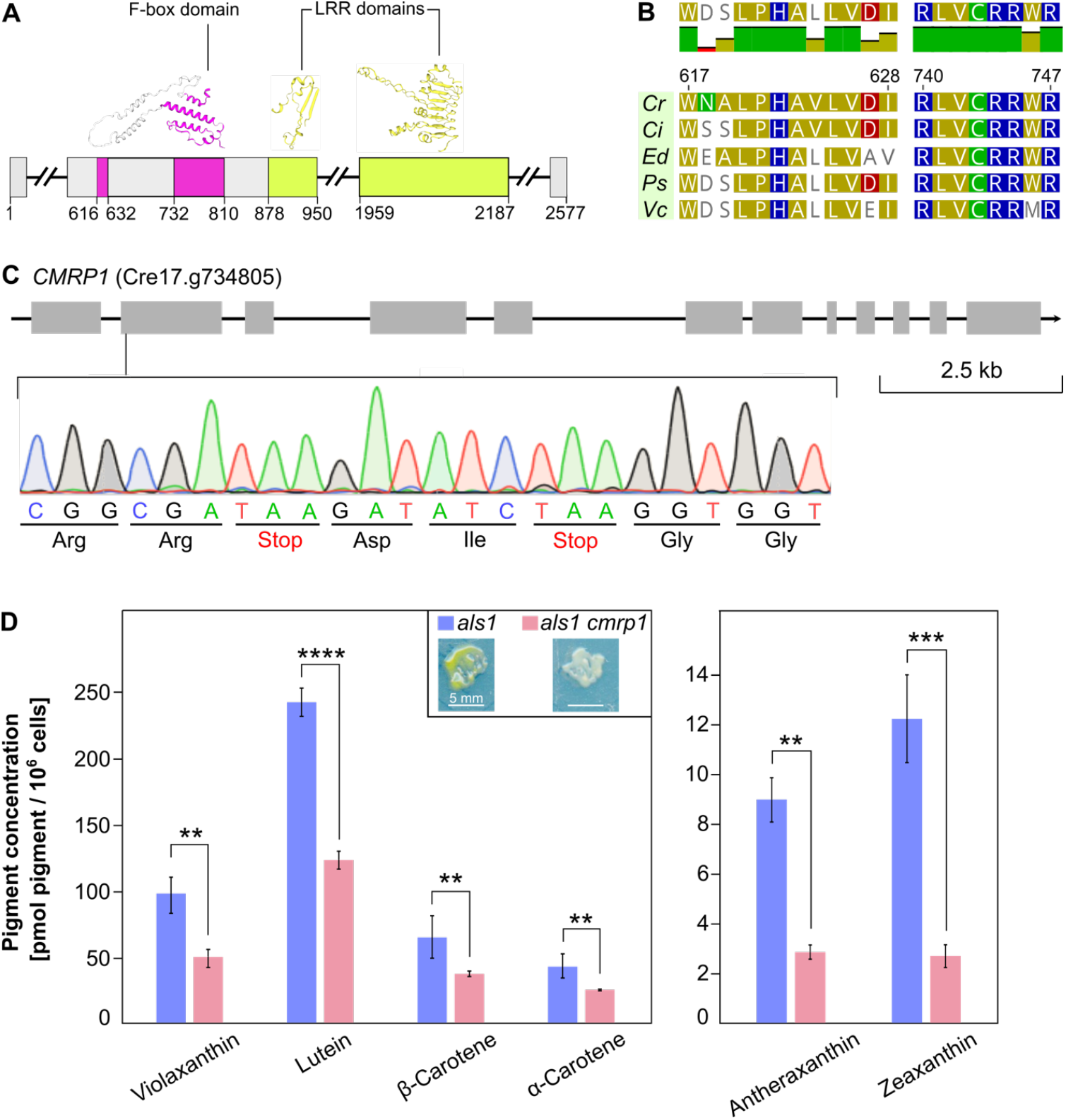
The gene *CMRP1* is involved in the regulation of carotenoid metabolism. **(A)** The predicted protein structure of CMRP1 (carotenoid metabolism-regulating protein). The leucine-rich repeat (LRR) domains are shown in yellow and the F-box domain in magenta. The three-dimensional structures were predicted by AlphaFold 3.0 (Thompson and Petrić Howe, 2024). **(B)** Conserved motifs in the CMRP1 F-box domain of volvocine algae. The motifs shown are highly conserved among eukaryotic F-box proteins and are involved in the formation of the SCF (Skp1-Cullin-F-box protein) ubiquitin ligase complex (Schulman et al., 2000). Residues identical to those of *C. reinhardtii* are highlighted by the same color. The consensus sequence is shown at the top of the alignment. Cr, *Chlamydomonas reinhardtii*; Ci, *Chlamydomonas incerta*; Ed, *Edaphochlamys debaryana*; Ps, *Pleodorina starrii*; Vc, *Volvox carteri*. **(C)** Construction of a *cmrp1* knockout mutant. Two in-frame stop codons were introduced in the 2^nd^ exon of *CMRP1* (Supplementary Method S1). CC-3840 was used as a background strain. **(D)** Carotenoid concentrations in the *als1* and *als1 cmrp1* strains under illumination with a light intensity of 50 µmol photons m^-2^ s^-1^. The *als1 cmrp1* mutant contains two nonsense mutations in the *CMRP1* gene (electropherogram in (C)) and a missense mutation in the *ALS1* gene. The strain *als1* contained only the *ALS1* mutation and was used for comparison. Only carotenoids whose concentrations were significantly different between the two strains are shown. The concentrations of the carotenoids neoxanthin and loroxanthin, and of chlorophylls *a* and *b* are shown in Supplementary Fig. S6. Statistical analysis is described in Fig. 2. The inset shows the color difference between the strains grown in the dark.

To verify whether *CMRP1* is involved in carotenoid metabolism in *C. reinhardtii*, we generated a knockout mutant by CRISPR/Cas9-mediated co-editing of *CMRP1* and the endogenous selectable marker gene *ALS1* (Supplementary Method S1). To insert in-frame stop codons, we manually revised the *CMRP1* gene model based on the RNA-seq data of OCM-43. As a result, we generated a knockout mutant with two stop codons in the *CMRP1* coding sequence (Fig. 4C). In contrast to OCM-43 with the upregulation of *CMRP1* and larger carotenoid amounts (Fig. 2A), the knockout mutant showed lower concentrations of the same pigments (Fig. 4D). This result confirms that *CMRP1* is involved in the regulation of carotenoid metabolism in *C. reinhardtii*, thus becoming the first example of a gene from green algae whose function has been uncovered with a gene overexpression screen.

## Discussion

The forward genetic approach, ERGO, used in our study is based on the random overexpression of genes in the *C. reinhardtii* nuclear genome. This method differs considerably from methods that are based on gene knockout. One of the main differences lies in the components of the transformation cassette used to generate the mutant library (Fig. 1A; Supplementary Fig. S2). Apart from the selectable marker gene, the cassette contains a DNA sequence capable of upregulating gene expression. One possibility is to use a strong promoter, as has been done in angiosperms (Memelink, 2003). However, promoters can only activate genes when inserted in the correct orientation directly upstream of the 5’-UTR (Andersson and Sandelin, 2020). In contrast, most enhancers can activate gene expression independently of their position, orientation, or distance with respect to the target gene (Schmitz et al., 2022; Marand et al., 2023) and are thus more advantageous for random gene overexpression. To date, the cauliflower mosaic virus (CaMV) 35S enhancer has been the most widely used enhancer for gene upregulation in angiosperms (Amack and Antunes, 2020), although only one example of moderate gene activation by this enhancer has been reported in *C. reinhardtii* (Ruecker et al., 2008). In this study, we used the previously characterized enhancer *E*_*hist cons*_ from the *C. reinhardtii* histone genes (Lihanova et al., 2024). The reason for this choice is the ability of *E*_*hist cons*_ to activate gene expression over a distance of at least 1.5 kb (Lihanova et al., 2024), making it especially advantageous for gene overexpression over larger genomic distances. Indeed, the most significantly upregulated gene in OCM-43, *CMRP1*, was activated over a distance of ca. 2 Mb (Fig. 3A). Although previous investigations in angiosperms showed that the integrated enhancer mostly upregulated proximal genes (Weigel et al., 2000), long-distance promoter-enhancer interactions are being increasingly reported in eukaryotes (Tjalsma et al., 2025; Bower et al., 2025). In a future study, chromatin conformation capture techniques (e.g. Sun et al., 2024), which have yet to be applied to *C. reinhardtii*, can be used to determine whether long-distance gene activation in OCM-43 was directly caused by *E*_*hist cons*_.

Another aspect to consider when using a forward genetic screen is the extent of changes in gene expression after enhancer integration. The RNA-seq data analysis of OCM-43 showed that insertion of an enhancer at a single location in the genome influences (directly or indirectly) over 2800 genes by up- or downregulating their expression (Fig. 3C; Supplementary Data Set 2). While some of these extensive changes may have been directly caused by the enhancer integration, it is also possible that enhancer-mediated gene activation influenced the expression of other genes, further increasing the number of differentially expressed genes in the whole genome. This scenario seems particularly plausible in OCM-43, where a gene for a putative F-box protein has been upregulated. Proteins from the F-box superfamily are major recognition subunits of the SCF (Skp1-Cullin-F-box protein) ubiquitin ligases (Zheng and Shabek, 2017). Their protein-interaction domains not only recognize the components of the ligase complex but also recruit the substrate proteins for proteasomal degradation (Hua et al., 2011). Moreover, F-box proteins can act as transcriptional co-activators (e.g. Rieu et al., 2023); if this scenario is true for OCM-43, it might be the reason why the transcription of so many genes is changed in the mutant compared to the background strain.

The final point for consideration when designing a forward genetic screen is the gene density in a genome. Compared to the model organisms with similar genome sizes such as *Arabidopsis thaliana, Drosophila melanogaster*, or *Caenorhabditis elegans, C. reinhardtii* has a substantially higher gene density of about 79%, which is caused mainly by the high abundance and length of introns (Craig and Vallon, 2023). Therefore, it is not surprising that 60% of the OCMs analyzed in the present study contained an enhancer insertion in an exon or intron, which likely resulted in loss of gene function (Supplementary Data Set 1). For instance, we isolated two OCMs with a cassette insertion in an open reading frame. The mutant OCM-12 was unable to accumulate lipids under nitrogen deficiency in the light due to the knockout of the *TAR1* protein kinase gene (Supplementary Fig. S7A, B), as already demonstrated in a previous study (Kajikawa et al., 2015). Another mutant, OCM-15, contained an enhancer inserted in the gene encoding subunit H of magnesium chelatase (Supplementary Fig. S7C), an enzyme involved in tetrapyrrole biosynthesis (Willows et al., 2023). As a consequence, this mutant was unable to produce chlorophyll in the light and exhibited a red-brown color, most likely caused by the accumulation of protoporphyrin IX and heme (Supplementary Fig. S7D). Although the phenotypes of OCM-12 and OCM-15 have been mainly caused by loss of gene function, an intragenically inserted enhancer may still have upregulated a proximal or a distal gene; this feature makes such mutants more challenging to characterize but does not eliminate their potential for studying gene function.

In conclusion, we introduced a forward genetic approach based on gene overexpression. We have successfully identified a gene with a previously unknown function, and further characterization of this gene will widen our knowledge on the biological roles of ubiquitous F-box proteins. In addition, the mutants generated in our study provide the scientific community with a useful tool to study the role of carotenoids in the mechanism of photoprotection. To our knowledge, this is the first time that random gene overexpression, previously only applied to angiosperms, has been successfully used in green algae, thereby opening up new possibilities for functional genomics research in the green lineage and beyond.

## Materials and Methods

### Generation of the *C. reinhardtii* mutant library

To generate a library of *C. reinhardtii* insertional mutants, the plasmid pAT_4x*E*_*hist cons*__*APHVIII* was designed (Supplementary Fig. S1). Its main components include enhancer *E*_*hist cons*_ and the *APHVIII* selection marker. The *E*_*hist cons*_ sequence was amplified from the plasmid pUC57-Amp (Lihanova et al., 2024) with the forward primer 5’-TGGCGGCCGCTCTAGAACTCTGACACATGCAGCTCC-3’ and reverse primer 5’-AGTTAACCGTACGTTCGAAGACCGTCATCACCGAAG-3’ (Eurofins Genomics, Ebersberg, Germany), and purified with the GenElute PCR Clean-Up Kit (Sigma-Aldrich, Merck, Burlington, MA, USA). To generate a plasmid backbone, the plasmid pOpt_mCerulean3_Paro with the *APHVIII* gene (Lauersen et al., 2015) was digested with *Bcu*I and *Bam*HI and purified with the GenElute Gel Extraction Kit (Sigma-Aldrich). For Gibson cloning (Gibson et al., 2009), 100 ng of plasmid backbone and a 3-fold molar excess of the *E*_*hist cons*_-containing PCR product were mixed with 10 µl of the NEBuilder HiFi DNA Assembly Master Mix (NEB, Ipswich, USA) in a final volume of 20 µl. After incubation at 50 °C for 1 h, 2 µl of the reaction mixture were used directly for the transformation of chemically competent *Escherichia coli* TOP10 cells using the heat shock method (Sambrook and Russell, 2006). For plasmid preparation, the GenElute Plasmid Miniprep Kit (Sigma-Aldrich) was used and the correct plasmid sequence was confirmed via Sanger sequencing (Genewiz, Azenta Life Sciences, Leipzig, Germany).

*C. reinhardtii* strain CC-3840 (Δ*chlL, spr-u-1-6-2, sr-u-2-60, mt*^*+*^; Suzuki and Bauer, 1992) was used for electroporation. To obtain between 100 and 300 single colonies per agar plate, the transformation procedure from Lihanova et al. 2024 was adjusted as follows. The PCR product containing *E*_*hist cons*_ and the *APHVIII* gene was amplified from the plasmid pAT_4x*E*_*hist cons*__*APHVIII* using a forward primer 5’-CTCTGACACATGCAGCTCC-3’, and a reverse primer 5’- ATGACCATGATTA?GCCAAGC-3’ (Eurofins Genomics). The resulting 2047 bp PCR product was purified with the GenElute PCR Clean-Up Kit (Sigma-Aldrich). For electroporation, the cell suspension of 1 × 10^8^ cells ml^-1^ was mixed with 100 ng DNA to favor one DNA insertion event per transformant (Gonzalez-Ballester et al., 2011). Electroporated samples were plated on agar plates containing 15 µg ml^-1^ paromomycin, and the plates were incubated at 20 °C in the dark for three weeks. The transformation efficiency for CC-3840 reached 3,700 ± 800 transformants µg^-1^ DNA. Colony PCR with the primers 5’-GTTGGATGGATGCGGCAGG-3’ and 5’- TATGTATAGCGGCAGAATAGTCG-3’ was used to check for the full-length insertion of *E*_*hist cons*_ in the nuclear genome.

### Pigment analysis with the high-performance liquid chromatography (HPLC)

10 ml of the liquid algal cultures, grown at 20 °C to a cell density of at least 1 × 10^6^ cells ml^-1^, was vacuum-filtered onto glass fiber filters (Type GF 6, Ø 125 mm; Carl Roth GmbH, product no. AY46.1) cut into smaller disks of Ø 10 mm. The filters were placed in a 7 ml Precellys tube along with a glass bead mixture (beads Ø 0.25-0.5 mm /beads Ø 0.75-1.0 mm = 3/1; Carl Roth, product no. A553.1, A554.1) and 2 ml of extraction solution (81% (v/v) methanol, 10% (v/v) ethyl acetate, 9% (v/v) 0.2 M ammonium acetate). Cells were lysed in a Precellys Evolution homogenizer (Bertin Technologies, Montigny-le-Bretonneux, France) with two rounds of 20 s at 6500 rpm and a break of 5 s in between. After disruption, the supernatant was centrifuged at 14,800 x g for 2 min. 100 μl of the supernatant were used for analysis with the Ultimate 3000 HPLC system equipped with a PDA-3000 photodiode array detector (Thermo Fisher Scientific, Waltham, Massachusetts, USA). The pigments were separated on a 250/4.6 Nucleosil 120-5 C18 reversed-phase column (WICOM GmbH, Heppenheim, Germany) at 25 °C using a constant flow rate of 1.2 ml min^-1^. The samples were eluted with a binary gradient consisting of 97.5% (v/v) acetonitrile + 2.5% (v/v) 25 mM Tris-HCl, pH 7.5 (eluent A), and 75% (v/v) methanol + 25% (v/v) ethyl acetate (eluent B). After 100% eluent A for 18 min, the gradient changed linearly to 100% eluent B over the course of 2 min, and was kept at 100% eluent B for another 15 min. The elution of pigments was monitored at 440 nm, and absorption spectra were recorded in the wavelength range from 400 nm to 750 nm during the separation. The pigments were quantified as described by Frommolt et al., 2001. For cultures grown in the dark, lutein was used as a standard instead of a chlorophyll *a* because *C. reinhardtii* CC-3840 does not produce chlorophyll *a* in the absence of light (Suzuki and Bauer, 1992). The de-epoxidation state (DEPS) of the xanthophyll cycle pigments was calculated with the formula:

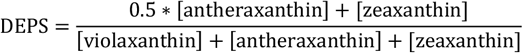

### Quantification of photosynthetic activity

To quantify photosynthetic activity, algal cultures were grown in liquid TP medium (Hui et al., 2022) at 20 °C under continuous white light (50 µmol photons m^-2^ s^-1^) and stirring (180 rpm) for three days. Before measurements, the cultures were diluted with the spent TP medium to a final cell concentration of 1 × 10^6^ cells/ml. For light stress experiments, the cultures grown at 50 µmol photons m^-2^ s^-1^ were illuminated with the light intensities of 1000 µmol photons m^-2^ s^-1^ or 2000 µmol photons m^-2^ s^-1^ for 4 h using an LED white light panel. The photosynthetic activity was quantified by measuring the quantum yield of PSII (Fv/Fm) using a chlorophyll fluorometer (Handy PEA+, Hansatech Instruments, King’s Lynn, United Kingdom). After dark-adapting the cultures for 5 min, a light pulse with an intensity of 3500 μmol photons m^-2^ s^-1^ was applied for 1 s. The high time resolution of the Handy PEA+ allowed the determination of the minimal fluorescence, Fo, at the beginning of the light pulse, whereas the maximum fluorescence, Fm, was detected at the end of the pulse. The variable fluorescence, Fv, was calculated as Fm-Fo.

### Whole-genome sequencing and mapping of enhancer insertion sites

For the sequencing of genomic DNA, liquid algal cultures were grown under continuous white light (50 µmol photons m^-2^ s^-1^) and shaking (150 rpm) for 5 days. After centrifugation at room temperature, 4000 x g for 5 min, 100 mg (fresh weight) of cell pellet were resuspended in 400 µl of the lysis buffer PL1 from the NucleoSpin Plant II Mini kit for DNA extraction from plants (Macherey-Nagel, Düren, Germany; product no. 740770). The resulting cell suspension was frozen in liquid nitrogen and thawed at 65 °C in a heat block with a total of four freeze-thaw cycles. The following steps were based on the manufacturer’s protocol. Genomic DNA was eluted by pipetting 25 µl of the preheated EB buffer (10 mM Tris-Cl, pH 8.5) on the center of the silica membrane, with incubation at 65 °C for 5 min and centrifugation at 11,000 x g for 1 min. This step was repeated and the two eluates combined. The DNA concentration and purity were assessed with a Nanodrop 2000/2000C Spectrophotometer (Thermo Fisher Scientific). Library preparation and paired-end Illumina sequencing were performed by Novogene (Munich, Germany) on the Illumina NovaSeq X Plus Series platform generating 1 GB of raw data per mutant with 150 bp paired-end reads and an average sequencing depth of 9-fold. For mapping of the enhancer insertion sites, the *C. reinhardtii* CC-4532 v6.1 reference genome (Craig et al., 2023) was indexed and the 150 bp reads were aligned to the reference genome using BWA-MEM v0.7.17 (Li, 2013). Discordant paired-end reads with one read mapping to the sequence of the transformation cassette and the other read mapping to a chromosomal location were filtered using SAMtools v1.6 (Li et al., 2009; Danecek et al., 2021). Filtered discordant reads were manually validated for each mutant using Integrated Genome Viewer (IGV v2.17.4).

For long-read sequencing, genomic DNA was extracted with the NucleoSpin Plant II Mini kit and eluted as described above using ultra-pure water instead of EB buffer. After elution, 50 µl of the Short Read Eliminator (SRE) buffer (SRE XS kit, PacBio of California Inc, CA, USA, product no. 102-208-200) were added to precipitate high-molecular-weight DNA. After centrifugation at 10,000 x g, 4 °C for 1 min, the colorless pellet was washed twice with 200 µl of 70% ethanol by centrifugation at 10,000 x g, 4 °C for 2 min each time. The DNA was air-dried at 20 °C and redissolved in 50 µl of EB buffer. Library preparation and single-molecule real-time sequencing (SMRT) were performed by Novogene on the PacBio Revio platform generating 1.1-1.4 GB of raw data per mutant with the mean read length ranging from 14,650 bp (OCM-26) to 16,854 bp (OCM-6). *E*_*hist cons*_ insertion sites were mapped with pbmm2 v1.2.0 by indexing the *C. reinhardtii* CC-4532 v6.1 reference genome and aligning the reads to the reference using the preset HIFI (https://github.com/PacificBiosciences/pbmm2). The resulting BAM files were sorted and indexed with SAMtools. Correct identification of the *E*_*hist cons*_ insertion sites for the chosen mutants was confirmed via Sanger sequencing (Genewiz). Insertion sites for all mutants are available in the Supplementary Data Set 1.

### RNA sequencing and gene expression analysis

For the mRNA sequencing, each liquid algal culture was grown in three biological replicates under continuous light (15 µmol photons m^-2^ s^-1^) and shaking (150 rpm) for 3 days. After centrifugation at room temperature at 3000 x g for 5 min, up to 100 mg (fresh weight) of cell pellet were resuspended in 350 µl of buffer RA1 containing 20 mM DTT (final concentration) from the NucleoSpin RNA Plant kit (Macherey-Nagel, product no. 740949). The resulting cell suspension was frozen in liquid nitrogen and thawed at 56 °C in a heat block with a total of four freeze-thaw cycles. The following steps were based on the manufacturer’s protocol. The RNA concentration and purity were assessed with a Nanodrop 2000/2000C Spectrophotometer (Thermo Fisher Scientific). mRNA library preparation and paired-end mRNA-seq were performed by Novogene on the Illumina NovaSeq X Plus Series platform generating 6 GB of raw data per mutant with 150 bp paired-end reads. For the gene expression analysis, the *C. reinhardtii* CC-4532 v6.1 reference genome was indexed with the parameter “--sjdbOverhang 149” and the reads were aligned to the indexed genome using STAR v2.7.11b (Dobin et al., 2013) with the parameter “--alignIntronMax 5000” to define the maximum intron length. The resulting BAM files were sorted and indexed with SAMtools. For differential gene expression analysis, the mapped reads were counted with htseq-count (Anders et al., 2015) and the generated count matrix was used as input for DESeq2 using the standard workflow (Love et al., 2014). To generate more accurate log_2_ fold change values, the log_2_ fold change estimates were shrunk towards zero (Zhu et al., 2019). Graphs were generated with R Statistical Software v4.2.1 and RStudio v2022.07.2 (R Core Team, 2021).

### Multiple sequence alignment and protein motif analysis

Homologous proteins, available as a multiple sequence alignment in Supplementary Data Set 3, were obtained by running four iterations of PSI-BLAST (Altschul et al., 1997) using the F-box core region with the insert sequence excluded. Multiple sequence alignment was performed using mafft v7.525 (Katoh and Standley, 2013) with the parameter “auto”. This alignment was used as input to predict structural motifs with HHpred available online as part of the MPI Bioinformatics Toolkit (Söding et al., 2005), using both the full alignment (no F-box detected) and an alignment with gappy columns removed (F-box detected). Alignment filtering was performed using trimAl v1.4.rev22 (Capella-Gutiérrez et al., 2009) and the parameter “-gt 0.2” to remove alignment columns with gaps in more than 80% of sequences (Supplementary Data Set 4).

### Generation of the knockout mutant by CRISPR/Cas9-mediated genome editing

The editing strategy was based on the co-editing of two genes: the *ALS1* selectable marker gene (Cre09.g386758), and the gene of interest, *CMRP1* (Cre17.g734805). Detailed protocol for design of crRNAs and homologous templates as well as a pipetting scheme for RNP assembly are available in Supplementary Method S1.

Electroporation of *C. reinhardtii* CC-3840 was based on the protocol from Lihanova et al. 2024, adjusted as follows. Liquid cultures, grown at 20°C under continuous white light (30 μmol photons m^-2^ s^-1^) and shaking (150 rpm) for 4 days, were diluted 3-fold and incubated for additional 16 h under the same conditions. After centrifugation at 1700 x g, 15 °C for 10 min, the cell pellet was washed twice with 10 ml of TAP medium containing 40 mM sucrose and 1 µg/ml cyanocobalamin and resuspended in the same medium to reach a final cell density of 2.2 × 10^8^ cells ml^-1^. 36 µl of the final cell suspension were mixed with a total of 4.5 µl of the ribonucleoprotein (RNP) complexes and homologous DNA templates. As a negative control, 4.5 µl of the IDT Duplex Buffer were added instead of the RNPs mixture. Electroporation was performed with the NEPA21 electroporator (Nepa Gene Co. Ltd., Ichikawa, Chiba, Japan) in precooled electroporation cuvettes with a 2 mm gap (EC-002S; Xceltis GmbH, Mannheim, Germany). The impedance was adjusted to 0.25-0.28 kΩ by adding more cell suspension to the cuvette if necessary. Two poring pulses with a starting voltage of 250 V were applied for 6 ms with a 50 ms pulse interval and a decay rate of 40%. The number of polarity-exchanged transfer pulses with a starting voltage of 20 V was set to five and applied for 50 ms with a 50 ms pulse interval and a decay rate of 40%. Directly after electroporation, the cell suspension was transferred from the cuvettes into 100 ml flasks with 20 ml of TAP medium containing 40 mM sucrose and 1 µg/ml cyanocobalamin. After incubation under white continuous light (10 µmol photons m^-2^ s^-1^) and shaking (100 rpm) for 3 h at 20 °C, samples were incubated with mild shaking (50 rpm) for 30 min at 39 °C, followed by recovery under white continuous light (10-15 µmol photons m^-2^ s^-1^) with shaking (100 rpm) for 40 h at 20 °C. For the plating of electroporated samples, 5 ml of TAP medium containing 0.5% (w/v) agar in a 15 ml conical centrifuge tube were preheated to 40 °C in a water bath. The cell suspension was centrifuged at 1,200 x g for 3 min. After centrifugation, the cell pellet was resuspended in 1 ml TAP medium, and 500 µl were gently mixed with 5 ml of the preheated TAP agar before plating on TAP plates containing 2% (w/v) agar and 5 µM sulfometuron methyl (SMM). The agar plates were incubated under continuous dim light (5-10 µmol photons m^-2^ s^-1^) for 3 days and then at a higher light intensity (50 µmol photons m^-2^ s^-1^) for another 7 days.

To verify gene editing, a colony PCR was pipetted using the primers 5’- CAGTGTGATCAAGGAGGCCTTTTAC-3’ and 5’-GACACTCCCCACCTCGATCTC-3’ for *ALS1*, and the primers 5’-CAGCTGGGGACAGTAGCCGCATTG-3’ and 5’-GTGTCGGCCGGAGAAGT GGCAG-3’ for *CMRP1*. Lysed cells from the SMM-resistant colonies were used as the PCR template. PCR products were analyzed by restriction digestion with *Eco*RV using 10 µl of the unpurified PCR product, 3 µl of the 10x digest buffer (Thermo Fisher Scientific), and 1 µl of the FastDigest restriction enzyme (Thermo Fisher Scientific) in a total volume of 30 µl. The mixture was incubated at 37 °C for 30 min and the reaction visualized via gel electrophoresis on a 1% (w/v) agarose gel in Tris-acetate-EDTA (TAE) buffer. PCR products digested by *Eco*RV were analyzed by Sanger sequencing (Genewiz).

## Supporting information

Supplementary Data Sets 1 and 2

Supplementary Data Sets 3 and 4

## Author contributions

S.S. conceived and supervised the project; Y.L., R.G., and S.S. designed experiments; Y.L. performed experiments and wrote the original manuscript draft; M.P. helped to generate the mutant library; R.J.C. helped with data analysis for mapping-by-sequencing, RNA-seq, generated multiple sequence alignments, and analyzed protein motifs; R.G. helped to quantify carotenoids and interpret the obtained data. All authors edited the manuscript.

## Supplementary Data

Supplementary Figure S1: Plasmid map of pAT_4x*E*_*hist cons*__*APHVIII*.

Supplementary Figure S2: Design of the enhancer-containing transformation cassette for ERGO in *C. reinhardtii*.

Supplementary Figure S3: Concentrations of neoxanthin, loroxanthin, and chlorophylls in OCM-43 and CC-3840.

Supplementary Figure S4: Verification of the *E*_*hist cons*_ sequence and insertion site in the OCM-43 nuclear genome by Sanger sequencing.

Supplementary Figure S5: Multiple sequence alignment of the F-box domain of CMRP1 homologs in the representative algae from Chlorophyceae and Trebouxiophyceae.

Supplementary Figure S6: Concentrations of neoxanthin, loroxanthin, and chlorophylls in the *als1* and *als1 cmrp1* strains.

Supplementary Figure S7: Knockout mutants OCM-12 and OCM-15 isolated during ERGO screen.

Supplementary Method S1: Generation of knockout mutants by CRISPR/Cas9-mediated genome editing.

Supplementary Data Set 1: Mapping of the *E*_*hist cons*_ insertion sites in the nuclear genome of *C. reinhardtii* orange-colored mutants (OCMs).

Supplementary Data Set 2: RNA-seq data analysis of OCM-43.

Supplementary Data Set 3: Multiple sequence alignment of CMRP1 homologs, unfiltered.

Supplementary Data Set 4: Multiple sequence alignment of CMRP1 homologs, filtered with trimAl.

