## Supplementary Figures for "Enhancer-driven random gene overexpression (ERGO): a method to study gene function in Chlamydomonas"

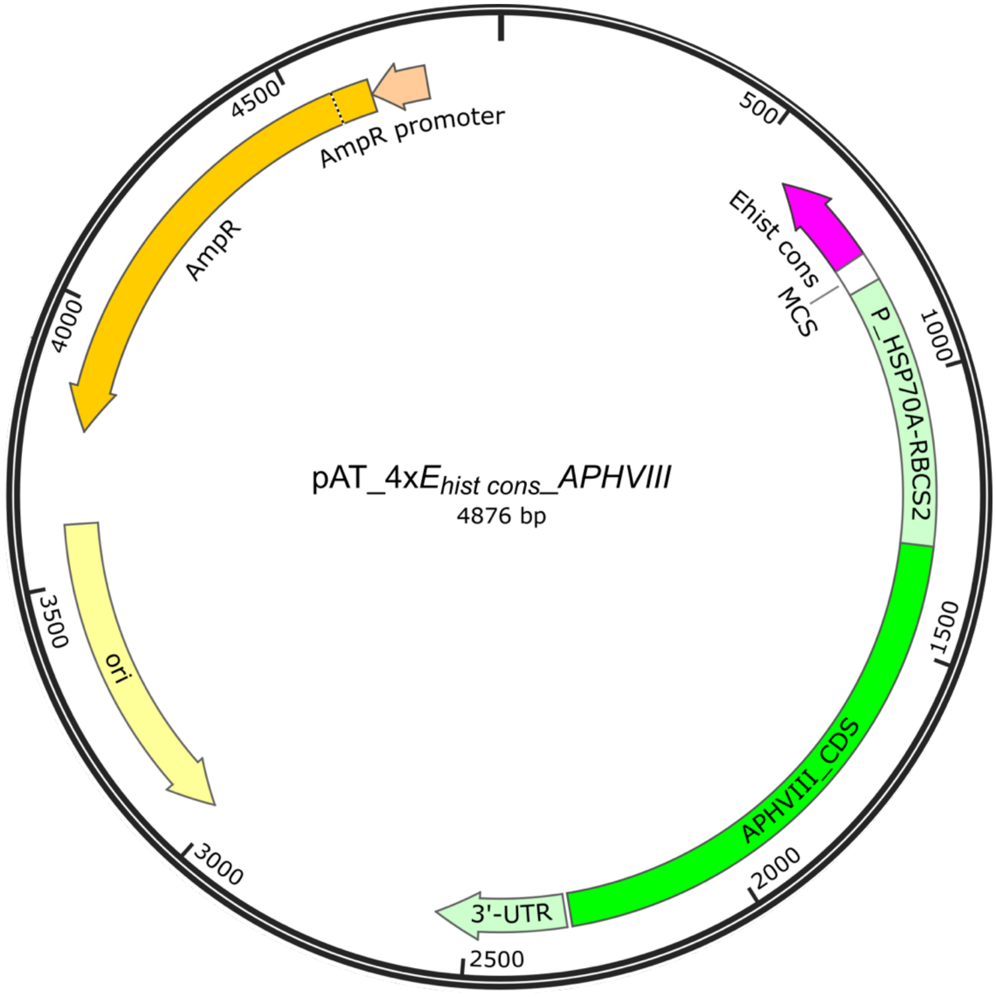


**Figure S1: Plasmid map of pAT_4x*E**hist cons*_*APHVIII*.** This plasmid contains the enhancer *E**hist cons* in the reverse orientation with respect to the *APHVIII* selection marker gene. The *APHVIII* gene under control of the tandem *HSP70A-RBCS2* promoter, the *RBCS2* 5'-UTR, and the *RBCS2* 3'-UTR originates from the pOpt_mCerulean3_Paro plasmid (Lauersen et al., 2015, Appl. Microbiol. Biotechnol. 99, 3491-3503). For selection of *E. coli* transformants and propagation of the plasmid in *E. coli*, the *bla* ($Amp^R$) gene and the origin of replication (*ori*) are included. MCS, multiple cloning site. The plasmid map was generated with SnapGene Viewer v5.3.2.

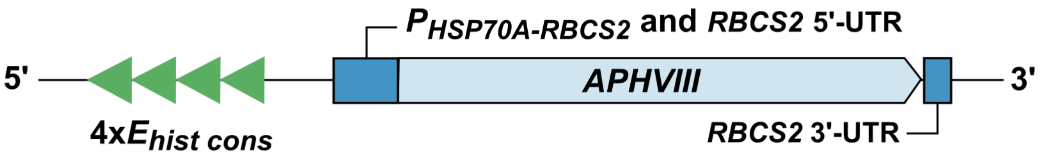


**Figure S2: Design of the enhancer-containing transformation cassette for ERGO in *C. reinhardtii*.** The *C. reinhardtii* nuclear genome was transformed with a 2047 bp PCR product containing the enhancer $E_{hist\ cons}$ with four copies of a conserved 8 bp-motif in the reverse orientation with respect to the *APHVIII* selection marker gene (four green triangles). The *APHVIII* selection marker under the control of the tandem *HSP70A-RBCS2* promoter, the *RBCS2* 5'-UTR, and the *RBCS2* 3'-UTR enabled transformant selection on paromomycin-containing agar plates. 50 bp flanking sequences on the 5'- and 3'-ends of the cassette originated from the plasmid pAT_4x$E_{hist\ cons}$_*APHVIII* backbone to protect $E_{hist\ cons}$ and the selection marker gene from truncation. UTR, untranslated region.

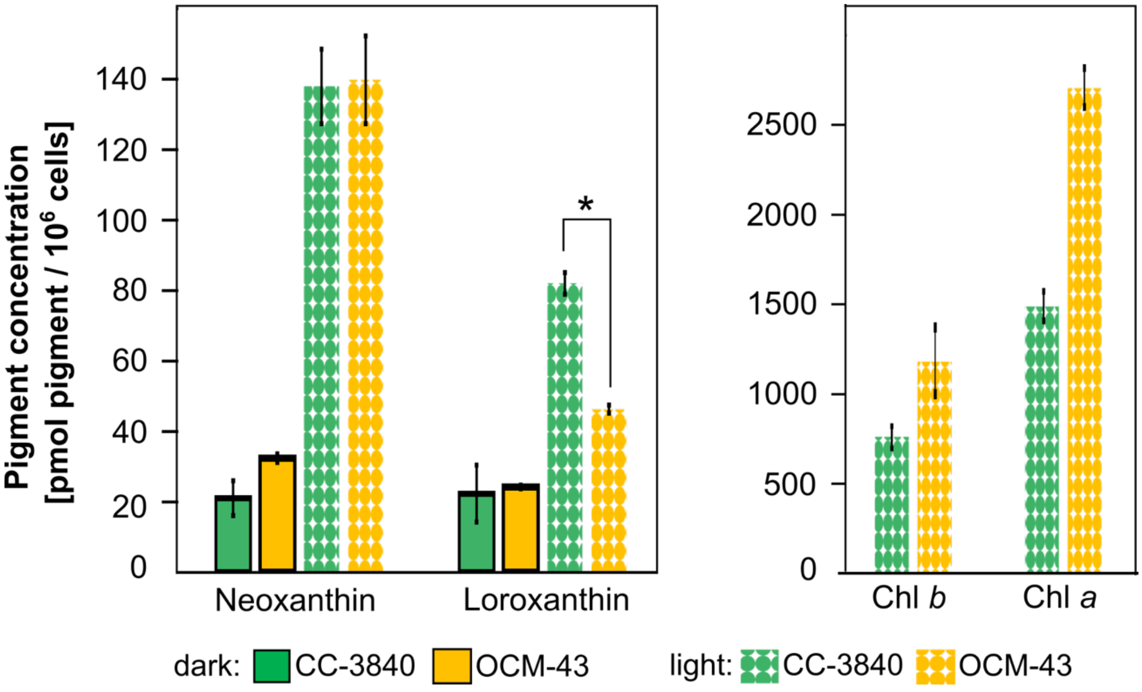


**Figure S3: Concentrations of neoxanthin, loroxanthin, and chlorophylls in OCM-43 and CC-3840.** TAP cultures were incubated in the dark or under illumination with a light intensity of 50 µmol photons $m^{-2}$ $s^{-1}$. Chlorophylls could only be produced in the light because of the mutation in the gene for the light-independent POR in the CC-3840 background strain (Suzuki and Bauer, 1992, Plant Cell 4, 929-940). Carotenoid concentrations were compared with each other using an unpaired t-test. Statistically significant differences between the mean values are indicated as follows: * $p \leq 0.05$. Data analysis was based on three biological replicates with the values indicating the mean ± standard deviation. Concentrations of other carotenoids are shown in Fig. 2A.

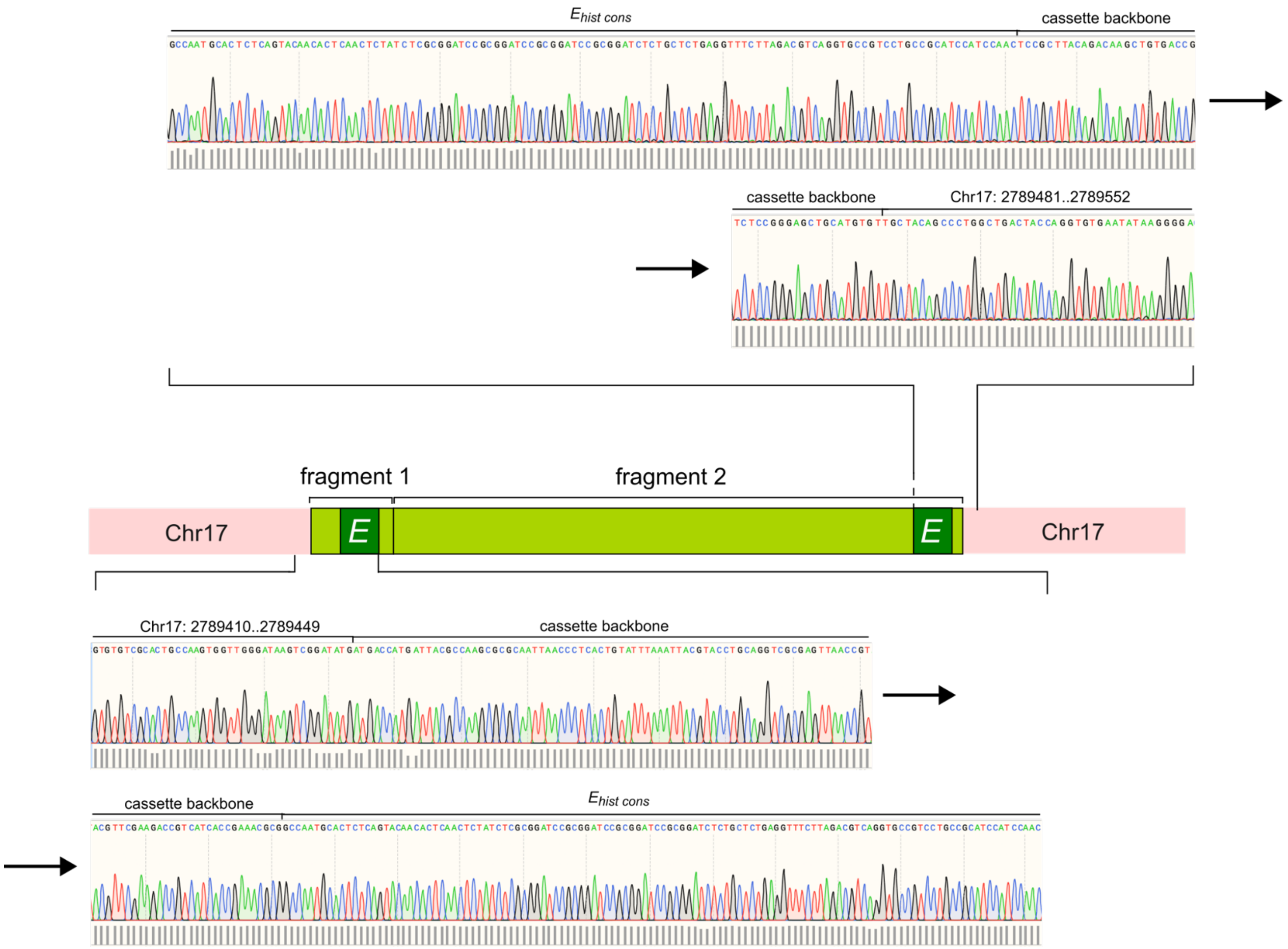


**Figure S4: Verification of the *$E_{hist\ cons}$* sequence and insertion site in the OCM-43 nuclear genome by Sanger sequencing.** Two fragments (fragment 1 and fragment 2) of the transformation cassette, shown as a light-green rectangle, were inserted on chromosome 17. $E_{hist\ cons}$ is depicted as a dark-green rectangle. Signal quality of electropherograms is represented by gray bars underneath each peak. Mapping-by-sequencing was based on the *C. reinhardtii* CC-4532 v6.1 genome (Craig et al., 2023, Plant Cell 35, 644-672). *E*, $E_{hist\ cons}$.

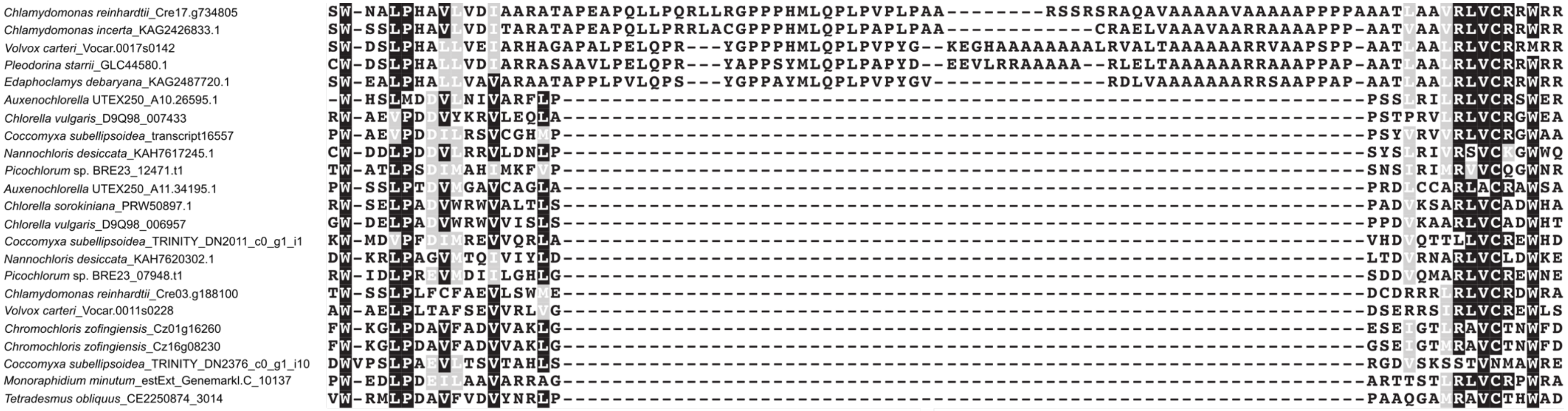


**Figure S5: Multiple sequence alignment of the F-box domain of CMRP1 homologs in the representative algae from Chlorophyceae and Trebouxiophyceae.** This alignment is based on the high-quality protein annotations. An insert of 99 amino acids is present only in volvocine algae. Motifs LPxxVLxxI and VRLVCRxW are highly conserved among eukaryotic F-box proteins and are involved in the formation of the SCF (Skp1-Cullin-F-box protein) ubiquitin ligase complex (Schulman et al., 2000, Nature 408, 381-386).

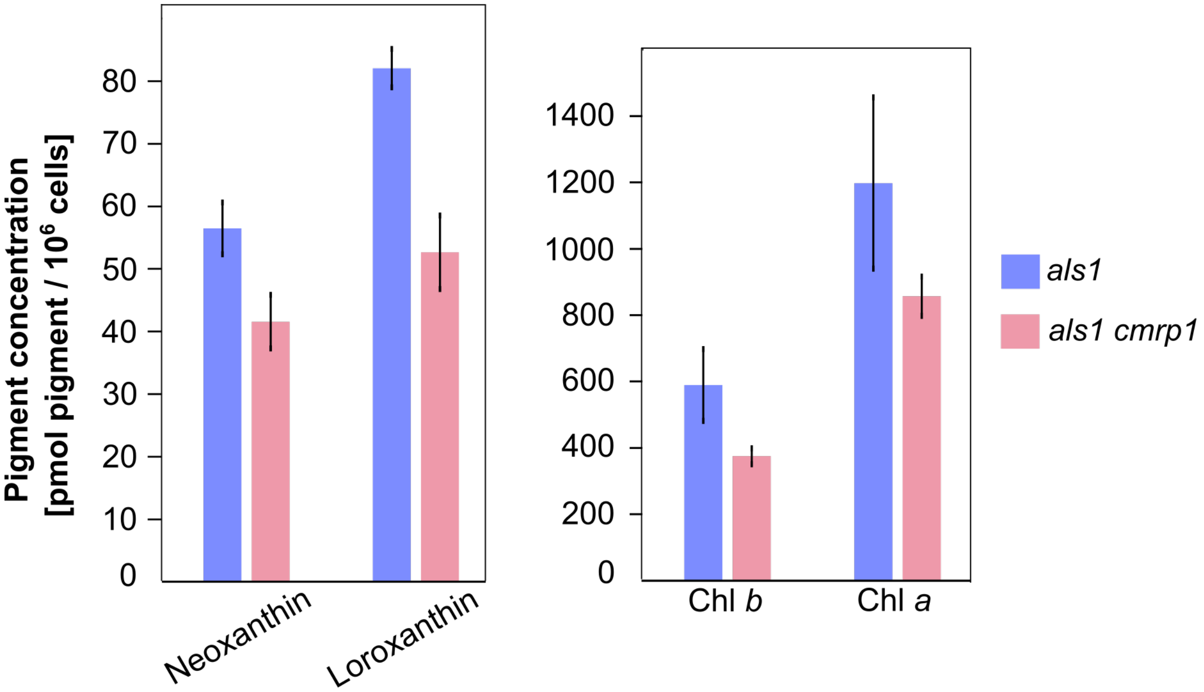


**Figure S6: Concentrations of neoxanthin, loroxanthin, and chlorophylls in the *als1* and *als1 cmrp1* strains.** Cultures were incubated under illumination with a light intensity of 50 µmol photons $m^{-2}$ $s^{-1}$. The strain *als1 cmrp1* contained a missense mutation in the *ALS1* gene and two nonsense mutations in the *CMRP1* gene. The strain *als1* contained only the *ALS1* mutation and was used for comparison as a reference strain. Carotenoid concentrations were compared with each other using an unpaired t-test; no statistically significant differences were observed between the two strains. Data analysis was based on three biological replicates with the values indicating the mean ± standard deviation. Concentrations of other carotenoids are shown in Fig. 4D.

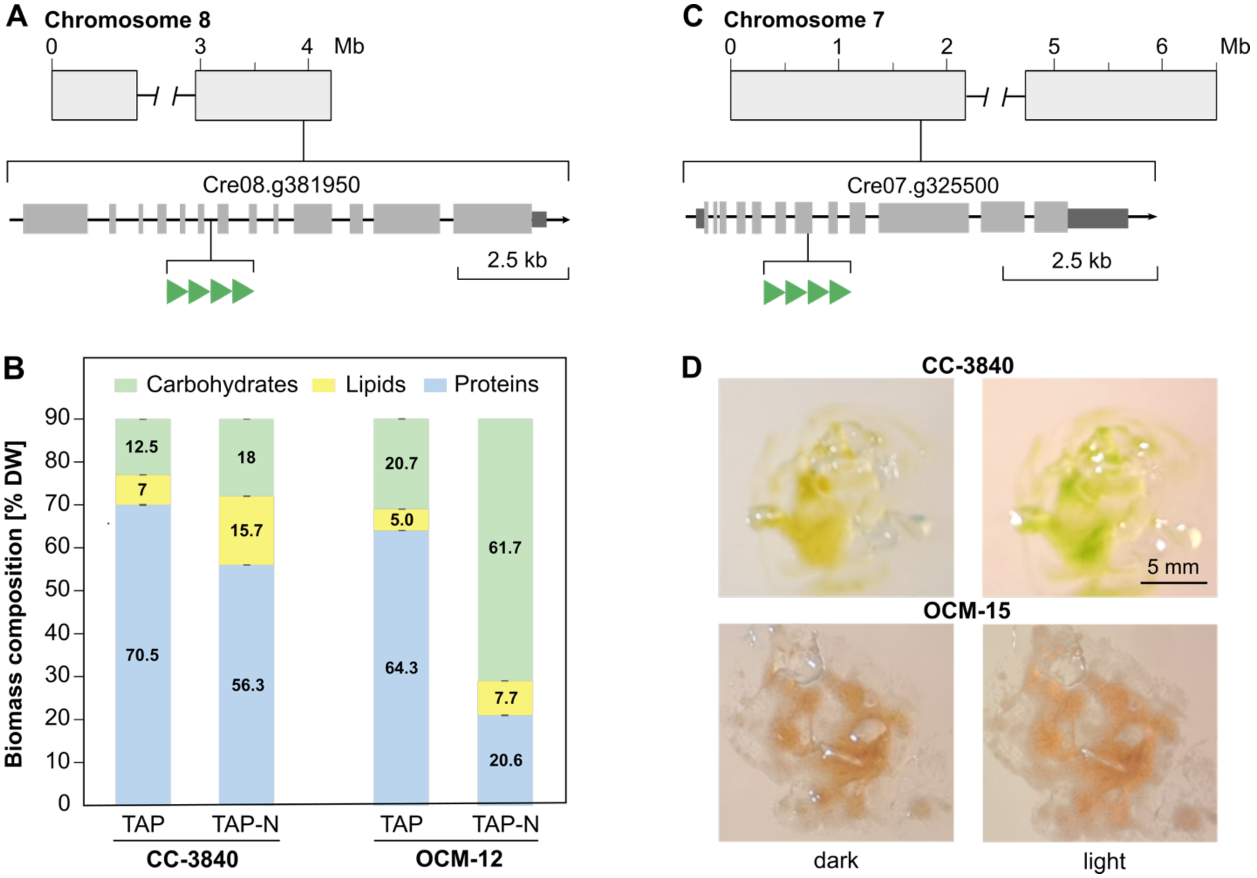


**Figure S7: Knockout mutants OCM-12 and OCM-15 isolated during ERGO screen. (A)** Insertion site of $E_{hist\ cons}$ in the OCM-12 nuclear genome. $E_{hist\ cons}$, shown by four green triangles, was inserted in the 6$^{th}$ intron of the gene Cre08.g381950 encoding a DYRK-type protein kinase. Gene model is based on the *C. reinhardtii* CC-4532, v6.1 genome (Craig et al., 2023, Plant Cell 35, 644-672). **(B)** Changes in carbohydrate, lipid and protein composition of OCM-12 and CC-3840 measured by FTIR (Fourier transform infrared spectroscopy). Liquid cultures were grown under continuous white light of 50 µmol photons $m^{-2}$ $s^{-1}$ in TAP medium or TAP-N medium (TAP medium without ammonium chloride) to reproduce the cultivation conditions from Kajikawa et al., 2015, Plant Phys. 168, 752-764. The FTIR measurements and data analysis were performed according to Wagner et al., 2010, J. Biophotonics 3, 557-566 with the assumption that proteins, carbohydrates and lipids make up 90% of the total dry weight. Data analysis was based on three biological replicates with the values indicating the mean percentage of dry weight (% DW) ± standard deviation. **(C)** Insertion site of $E_{hist\ cons}$ in the OCM-15 nuclear genome. $E_{hist\ cons}$, shown by four green triangles, was inserted in the 7$^{th}$ exon of the gene Cre07.g325500 encoding the subunit H of the enzyme Mg-chelatase. Gene model is based on the *C. reinhardtii* CC-4532, v6.1 genome (Craig et al., 2023). **(D)** Light-induced chlorophyll biosynthesis of CC-3840 compared to OCM-15. The cells were cultivated on a TAP agar plate in the dark or under continuous white light (50 µmol photons $m^{-2}$ $s^{-1}$) for 20 h. Compared to CC-3840, OCM-15 did not show a color change to green because of its inability to produce chlorophyll in the light.
