## Supplementary Method S1 for "Enhancer-driven random gene overexpression (ERGO): a method to study gene function in Chlamydomonas"

### Generation of knockout mutants by CRISPR/Cas9-mediated genome editing

This method is based on Akella et al., 2021, *Plant Physiology* 187, 2637-2655, where the gene *ALS1* (Cre09.g386758) was introduced as a candidate selection marker for future studies. Our method employes co-editing of *ALS1* and a gene of interest (in our case the *CMRP1* gene, Cre17.g734805) for the first time.

#### Design of crRNAs and homologous DNA templates

crRNAs were designed using the Cas-Designer (www.rgenome.net/cas-designer) with the following parameters:

- PAM type: *Sp*Cas9 from *Streptococcus pyogenes* (5'-NGG-3')
- Target genome: *Chlamydomonas reinhardtii* (v5.0)
- GC-content: 40-60%
- Out-of-frame score: at least 66
- Mismatches: 1 – 0 – 0

The best crRNA candidates based on the Cas-Designer prediction were synthesized as custom chemically modified 36 nt sequences with 16 additional nucleotides for annealing to the Alt-R® CRISPR-Cas9 tracrRNA (IDT; product no. 1072532) by the Integrated DNA Technologies (IDT; San Diego, CA, USA). The single-stranded oligodeoxynucleotides (ssODNs) were used as homologous templates and synthesized as Ultramer DNA Oligos (IDT) overlapping the Cas9 cleavage site with equal length of homology arms (at least 40 nt on both sides) and total length of 80-120 nt. The ssODN sequence for targeting of the gene of interest should contain at least one in-frame stop codon 5'-TAA-3' and an *Eco*RV recognition site 5'-GATATC-3' for PCR-based screening of transformants (see Materials and Methods for details). Sequences of crRNAs and ssODNs, used in this study, are summarized in Table S1.

**Table S1: Sequences of designed crRNAs and single-stranded homologous templates (ssODNs).** The displayed crRNA sequences anneal to the complementary target DNA region, with each crRNA additionally containing a 16 nt common sequence (GUUUUAGAGCUAUGCU) at its 3'-end for annealing to tracrRNA. Changes in the sequence of single-stranded DNA templates are written in lower case and highlighted in red. Asterisks at 5'- and 3'-ends of the templates denote for the phosphorothioate bonds, which inhibit the exonucleolytic degradation. crRNA and ssODN for the *ALS1* gene originated from Akella et al., 2021.

| | |
|---|---|
| **crRNAs** (5'-3') | |
| *ALS1*, exon 8 | GCUGCUGCUGGAUGUCCUU |
| *CMRP1*, exon 2 | GACCCGUUUCGCUGCAAGAG |
| **ssODNs** (5'-3') | |
| *ALS1*, exon 8 | C*C*G*CACCGGCCGGCCCGGCCCTGTGCTGGTGGACGTGCCCA**c**GGA**t**ATCCAGCAGCA GCTGGCGGTGCCGGACTGGGAGG*C*G*C |
| *CMRP1*, exon 2 | G*C*T*GTCACCGGCGCCGCCGCCACCACC**ttag**AT**at**CT**ta**TCGCCGCTCTTGCAGCGAAAC GGGTCTTGCCTGTTGCGGTGGTGGGGA*T*C*G |

**Pipetting scheme for RNPs and ssODN-mix (with example)**

- for 4 samples (half of each sample will be plated on a TAP-SMM (5 µM) agar plate resulting in 8 agar plates in total, with at least 50 colonies per plate to easily distinguish and isolate single colonies)

Pipette the following in 0.2 µl PCR-tubes:

| RNP1 (*ALS1*) | | RNP2 (gene of interest) | |
|---|---|---|---|
| **crRNA** | **tracrRNA** | **crRNA** | **tracrRNA** |
| Dilution of the stock: 2 nmol RNA + 13.3 µl $dH_2O$ -> ca. 150 µM | Dilution of the stock: 5 nmol RNA + 33.3 µl $dH_2O$ -> ca. 150 µM | Dilution of the stock: 2 nmol RNA + 13.3 µl $dH_2O$ -> ca. 150 µM | Dilution of the stock: 5 nmol RNA + 33.3 µl $dH_2O$ -> ca. 150 µM |
| 1.36 µl + 1.36 µl | | 4.1 µl + 4.1 µl | |
| + 1.13 µl Duplex Buffer = 3.85 µl RNA-Duplex (53 µM final conc. of each RNA) | | + 3.4 µl Duplex Buffer = 11.6 µl RNA-Duplex (53 µM final conc. of each RNA) | |
| PCR cycler: 96°C, 3 min – cool down slowly (take out of cycler and leave at room temperature) | | PCR cycler: 96°C, 3 min – cool down slowly (take out of cycler and leave at room temperature) | |
| + 1.1 µl Cas9 directly to the RNA-Duplex (RNA-Duplex [3.85 µl] : Cas9 [1.1 µl] = 3.5 : 1) = <u>ca. 5 µl RNP1</u> | | + 3.3 µl Cas9 directly to the RNA-Duplex (RNA-Duplex [11.6 µl] : Cas9 [3.3 µl] = 3.5 : 1) = <u>ca. 15 µl RNP2</u> | |
| PCR cycler: 37°C, 20 min – use for MasterMix (see below), or store at -80°C | | PCR cycler: 37°C, 20 min – use for MasterMix (see below), or store at -80°C | |

| **MasterMix, 5x** | | |
|---|---|---|
| RNP1 : RNP2 = 1 : 3 | | |
| 3.75 µl RNP1 (0.75 µl per sample) | + | 11.25 µl RNP2 (2.25 µl per sample) |
| + 7.5 µl ssODN-MasterMix * (1.5 µl per sample) | | |
| = <u>22.5 µl total</u> (4.5 µl per sample) | | |

*** ssODN-MasterMix:**

3.33 µl of 150 µM *ALS1*-ssODN (50 µM final concentration)
+ 5 µl of 300 µM of ssODN of gene of interest (150 µM final concentration)
+ 1.67 µl $dH_2O$
= <u>10 µl total</u>
